# Late consolidation of rRNA structure during co-transcriptional assembly in *E. coli* by time-resolved DMS footprinting

**DOI:** 10.1101/2024.01.10.574868

**Authors:** Yumeng Hao, Ryan M. Hulscher, Boris Zinshteyn, Sarah A. Woodson

## Abstract

The production of new ribosomes requires proper folding of the rRNA and the addition of more than 50 ribosomal proteins. The structures of some assembly intermediates have been determined by cryo-electron microscopy, yet these structures do not provide information on the folding dynamics of the rRNA. To visualize the changes in rRNA structure during ribosome assembly in *E. coli* cells, transcripts were pulse-labeled with 4-thiouridine and the structure of newly made rRNA probed at various times by dimethyl sulfate modification and mutational profiling sequencing (4U-DMS-MaPseq). The in-cell DMS modification patterns revealed that many long-range rRNA tertiary interactions and protein binding sites through the 16S and 23S rRNA remain partially unfolded 1.5 min after transcription. By contrast, the active sites were continually shielded from DMS modification, suggesting that these critical regions are guarded by cellular factors throughout assembly. Later, bases near the peptidyl tRNA site exhibited specific rearrangements consistent with the binding and release of assembly factors. Time-dependent structure-probing in cells suggests that many tertiary interactions throughout the new ribosomal subunits remain mobile or unfolded until the late stages of subunit maturation.

## Introduction

During active growth, *E. coli* cells synthesize more than 3,000 ribosomes per minute, in a process that requires the proper folding of the long pre-rRNA during transcription and the addition of ∼50 ribosomal proteins (r-proteins) to the small and large subunits (1). Although assembly is remarkably efficient and accurate in *E. coli* (2), variable folding of the pre-rRNA during transcription can delay the stable recruitment of ribosomal proteins until multiple proteins within a domain are able to bind the transcript (3,4). This raises the question of how the cellular environment and ribosome assembly factors overcome this delay, so that most pre-rRNA transcripts assemble within a few minutes. In bacteria, 10-30 assembly factors facilitate maturation by temporarily stabilizing native interactions (5-7) or preventing premature docking of incompletely folded domains (8). The assembly factors also prevent immature subunits from initiating protein synthesis (8-10).

The kinetics of rRNA folding have been difficult to probe *in situ* because assembly intermediates represent only a few percent of all ribosomes in cells. Previous studies have increased the numbers of immature particles by depleting cells of r-proteins or specific assembly factors. However, such mutations can reroute ribosome biogenesis through new assembly pathways (5,11-13), making it difficult to know which steps of rRNA folding are important for ribosome biosynthesis during normal growth. Affinity purification of assembly factor complexes can enrich for intermediates, but this approach often captures later complexes that are more stable (14). Here, we show that metabolic labeling of newly made rRNA, combined with time-resolved DMS footprinting, can reveal the slow folding dynamics of the rRNA in cells.

Reconstitution of the *E. coli* 30S and 50S ribosomal subunits *in vitro* (15-18) revealed a hierarchy of ribosomal protein (r-protein) addition and coupled conformational changes in the rRNA that occur as the components of the ribosomal subunits come together (19-21). Metabolic labeling of r-proteins with quantitative mass spectrometry and cryo-electron microscopy of partial ribosomal complexes from *E. coli* cells showed that this hierarchy is preserved in cells (22), with assembly following the 5’ to 3’ path of transcription, as expected (23). The mRNA decoding center in the 30S subunit and the peptidyl transfer center (PTC) in the 50S subunit are among the last regions to fold.

Despite these important advances, regions of the rRNA that fold or change conformation during assembly are typically too flexible to be resolved by cryo-EM and are invisible to methods that report protein binding. Chemical footprinting methods, however, can reveal changes in rRNA structure that occur during assembly. For example, in-cell hydroxyl radical footprinting of immature pre-30S ribosomes in the absence of assembly factors RbfA or RimM showed that 16S rRNA in the decoding site was unfolded and exposed to solvent (8). Similarly, *in vitro* and in-cell modification by dimethyl sulfate (DMS) and kethoxal revealed perturbations to the rRNA structure in immature ribosomes (5,12,24,25). DMS preferentially methylates the Hoogsteen face of exposed guanosines and the Watson-Crick face of unpaired adenosines and cytosines. When combined with high-throughput sequencing, DMS modification can capture structural perturbations across the transcriptome (26-28).

To probe the conformations of pre-ribosomes in real time in logarithmically growing *E. coli* cells, we combined DMS-MaPseq footprinting (29) with metabolic labeling of newly made rRNA with a 4-thio uridine analog (^4S^U) (30-32). We previously showed that ^4S^U labeling allows for selective analysis of newly transcribed rRNA in *E. coli* under conditions that minimally perturb cell growth (33). Because DMS readily penetrates cell membranes, the cellular RNA is methylated within 20-30 s, resulting in an overall 60-90 s measurement time that can capture slow steps of ribosome assembly and subunit maturation. The results reveal widespread remodeling of primary r-protein binding sites and nucleotides around the mRNA binding site before new ribosomes enter the 70S pool.

## METHODS

### Reagents, biological resources and computational resources

The sources for key reagents are indicated in the appropriate sections below and listed in Table S1. Oligonucleotides used in this study are listed in Table S2.

### Polysome profile and tritium labeling

*E. coli* MRE600 cells (*rna*^−^) were pulse-labeled with ^3^H-uridine for 0-2 min and harvested at various times after treatment of cells with rifampicin or chased with uridine. Samples were collected at various times after rifampicin addition or uridine addition, and polysome profiles were obtained as previously described (8). See Supplementary Information for additional detail.

### In-cell DMS modification of mature ribosomes

5 mL of LB media were inoculated with a single MRE600 colony and grown overnight at 37 °C with shaking. The following day, the culture was diluted at 1:1000 into 5 mL LB media and was grown at 37 °C to OD_600_ = 0.6. For in-cell dimethylsulfate (DMS) modification, the DMS solution was freshly prepared by diluting the DMS stock (cat. 41610, Sigma, USA) 1:3 with 100% ethanol. 5 mL diluted DMS or 5 mL water (no dose control) was added to 50 mL culture and mixed vigorously for 30 s before adding to 50 mL ice-cold quench buffer containing 50% 1.4 M 2-mercaptoethanol and 50% isoamyl alcohol before harvesting cells at 5,000 × *g* for 10 min. The pellets were washed with 50 mL 0.7 M 2-mercaptoethanol and harvested twice more and stored at –80 °C before RNA extraction.

### 4-thiouridine labeling and *in vivo* DMS modification of nascent transcripts

5 mL LB media were inoculated with a single MRE600 colony and grown overnight at 37 °C with shaking. The following day, the culture was diluted at 1:1000 into 500 mL LB media and was grown at 37 °C to OD_600_ = 0.6. For rifampicin chase experiments, 4-thiouridine (0.5 mM final) (cat. T4509, Sigma, USA) was added to 500 mL culture with thorough shaking. After 1 min, rifampicin was added to a final concentration of 200 µg/mL, and 50 mL aliquots of culture were removed for DMS modification at 0, 0.5, 1, 2, 3, 5 and 10 min after rifampicin addition (DMS samples with rifampicin treatment, N=7). Also, 50-mL aliquots were collected at 0, 6 and 10 min after rifampicin addition for no dose controls (No dose controls with rifampicin treatment, N=3). The samples were immediately modified with DMS and quenched with 2-mercaptoethanol as described above. For ^4S^U control samples (DMS controls, N=2), 0.5 mM 4-thiouridine was added to 50 mL culture with thorough shaking. After 10 min, the samples were treated with DMS as described above.

### RNA isolation

Total RNA was isolated from bacterial cells using the RNeasy mini kit (5 mL culture) and RNeasy maxi kit (50 mL culture) (Qiagen). RNA concentrations were measured using a Nanodrop spectrometer (ThermoScientific). ^4S^U-labeled RNA (100 µg) was biotinylated with HPDP-biotin (cat. 21341, ThermoScientific) and captured on 150 µL of streptavidin magnetic beads (NEB) per RNA sample. The recovered ^4S^U labeled RNA was about 1% of the total RNA. See Supplementary Information for further details.

### DMS-MaPseq sample preparation

Library samples for high-throughput sequencing and mutational profiling were prepared as previously described (26) and as outlined in detail in Supplementary Information. In brief, after fragmentation of the RNA and adaptor ligation, the RNA was copied with a thermostable group II-intron reverse transcriptase (TGI-RT III; InGex, USA). The resulting cDNA products were circularized with CircLigase ssDNA ligase (Lucigen, USA). Illumina sequencing adaptors were introduced by PCR and the final library sequenced at the JHU Genetic Resources Core Facility on a HiSeq2500 (Illumina) in Rapid Mode using 50 bp single reads. The reads were processed and aligned to the consensus sequence of the *rrn* operon (see Supplementary Information).

### Data analysis

#### Modification of mature ribosomes

The mutation rates in DMS-modified samples were compared to the corresponding mutation rate in no dose (ND) samples, for each nucleotide. A and C residues with mutation rates ≥ 2 fold higher in the DMS-treated samples than ND samples were clustered based on their mutation rates (see Fig. S2 and S3A). Alternatively, the diffBUM-HMM pipeline was used to analyze the modification pattern as previously described (34). Two sets of untreated (ND) and DMS-treated samples were used for each analysis. The read depths and mutation counts for each nucleotide were used to calculated posterior p values. Nucleotides that had UM values larger than 0.95 were selected as nucleotides that have high mutation rates and are presented in Fig. S3B.

#### Relative mutation ratios

To compare new and old ribosomes, DMS modification of ^4S^U-labeled RNA at various times after rifampicin addition was compared to DMS modification of the mature reference sample. For each nucleotide, we calculated the ratio and average of the mutation rates in the new rRNA and mature reference RNA, and defined the ratio as relative mutation ratios (RMR):

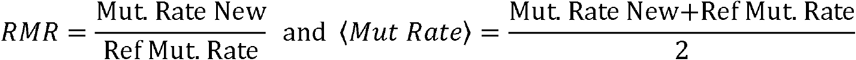

Nucleotides with average mutation rates corresponding to the bottom 10 percentile of values corresponded to helical residues that are base paired in both samples (233 nt total) and were excluded from further analyses. Nucleotides within the top 2 percentile of average mutation rates (36 nt total) mainly correlate with natural modifications that result in a very high mutation rate in both DMS-treated samples and ND controls. These residues were considered separately.

The sample treated with DMS 1 min after rifampicin had the lowest median RMR and represented the state that was the closest to the mature ribosome. The median RMR of this 1 min post-rifampicin sample, 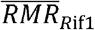, was used to benchmark the perturbations to new ribosomes for the entire time course. Nucleotides with 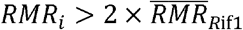 were designated ‘enhanced’, and those with 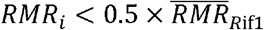 were designated ‘protected’. The enhanced and protected nucleotides were mapped on the rRNA secondary structures with xRNA and 3D structures with PyMOL.

#### Kinetic clusters

The enhanced and protected nucleotides were further clustered based on their change in DMS accessibility over time after rifampicin treatment as judged by the difference in RMR between two adjacent time points. Nucleotides with the largest change over time (top 50 percentile of standard deviation values, in which the standard deviation was calculated among the RMR values at different times) were clustered as listed in Table S3 and Fig. 4. K-means and other clustering methods were not more successful owing to the spread of MR values. See Table S4 for a list of residues in each cluster.

#### Solvent exposed bases

The solvent exposed surface areas for A N1 and C N3 atoms in the 70S ribosomes were calculated from pdb: 4ybb (35) structure in PyMOL using the get_area command with dot_solvent=1, dot_density=3, solvent_radius=3.

## RESULTS

### Time-resolved 4-thiouridine DMS footprinting (4U-DMS-MaP)

To monitor structural changes in the rRNA at different intervals after transcription, we combined 4-thiouridine (^4S^U) labeling with in-cell DMS footprinting (Fig. 1A). We first optimized the pulse labeling protocol and measured the overall ribosome assembly kinetics using ^3^H-uridine and sucrose gradient sedimentation (Fig. S1). We obtained the most uridine incorporation in the rRNA after 1 min (Fig. S1A,B). This labeling period was used for all subsequent experiments. When cells were treated with rifampicin to inhibit subsequent transcription initiation, the ^3^H-U labeled rRNA was mainly distributed among light fractions corresponding to 30S and 50S subunit assembly intermediates (Fig. S1C). Labeling peaked at 60 s for 30S subunits and 120 s for 50S subunits, but then increased again at 4-5 min (Fig. 1B and Fig. S1D), finally entering the 70S pool at 5-10 min (Fig. S1E-F). This pattern suggests that we are capturing two populations of subunits with different maturation kinetics, with the later population possibly reflecting the accumulation of stalled intermediates. For simplicity, we confined our footprinting analysis to the first two minutes after pulse-labeling, which is comparable to previous estimates of assembly kinetics in *E. coli* (2,36).

**Fig 1.**
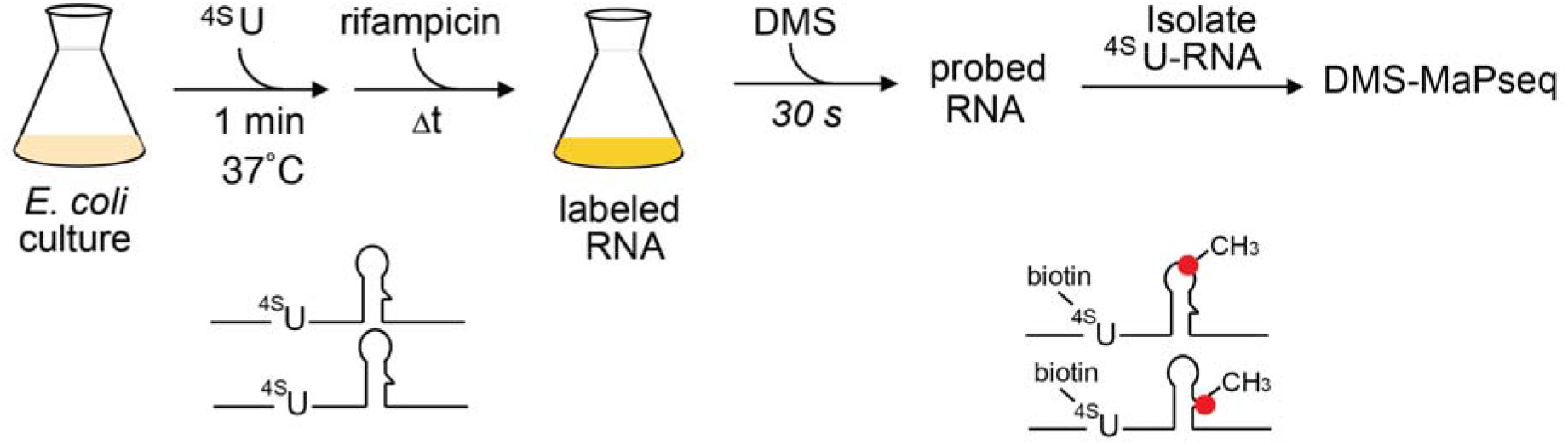
Visualizing the structures of newly made ribosomes with 4U-DMS-MaP. A. Overview of metabolic labeling of new transcripts and DMS footprinting in E. coli. Labeling with 4-thiouridine was stopped by rifampicin to inhibit transcription initiation or chased with excess uridine; 1 min ^4S^U labeling and 30 s of DMS treatment were optimal for high-throughput footprinting of new rRNA (Fig. S1A). B. Assembly kinetics based on the amount of ^3^H-uridine in peak fractions of sucrose gradient polysome profiles (Fig. S1D, H). The RNA was labeled with ^3^H-uridine for 1 min at 37 ºC before rifampicin addition (solid lines), or for 2 min at 25 ºC before uridine addition (dashed lines). The ^3^H-U content in the 30S and 50S peaks are plotted in red and blue, respectively. The labeled subunits accumulate about twice as slowly at 25 °C than at 37 °C.

For DMS footprinting, total RNA in logarithmically growing *E. coli* MRE600 cultures was pulse-labeled with 4-thiouridine (^4S^U) for 1 min before the addition of rifampicin. At four intervals from 0 to 2 min after rifampicin addition or a uridine chase, the cells were treated with DMS for 30 s, so that the shortest measurement time was 0.5 min post-labeling, and the longest time was 2.5 min post-labeling (Fig. 1). After DMS treatment, the ^4S^U-labeled RNA was biotinylated, isolated on streptavidin beads (37), and sequenced by high-throughput mutational profiling of DMS modified bases (29,38).

As expected, methylated A’s and C’s from DMS treated cells had 2 to 436-fold higher mutation rates than untreated (ND) samples, whereas the mutation rates of G’s and U’s were much less affected by DMS (Fig. 2A and Fig. S2). The reference DMS modification pattern of unlabeled mature ribosomes correlated well with the known secondary structures of the 16S, 23S and 5S rRNA (Fig. S3A) and the solvent accessibility of individual bases computed from structures of the mature ribosome (see Methods and (39)). A similar set of modified residues was identified by a recent hidden Markov algorithm for structure probing data (diffBUMM; (34), reflecting the robust signal-to-noise ratio of our data and the similarity of biological replicates (Fig. S3B). Control experiments in which *E. coli* cells were labeled with ^4S^U for 10 min before treatment with DMS yielded a very similar DMS modification pattern for the isolated ^4S^U-rRNA and unlabeled rRNA, showing that ^4S^U-labeling neither biases the DMS modification pattern nor appreciably interferes with ribosome assembly (Fig. S3C, D).

**Fig 2.**
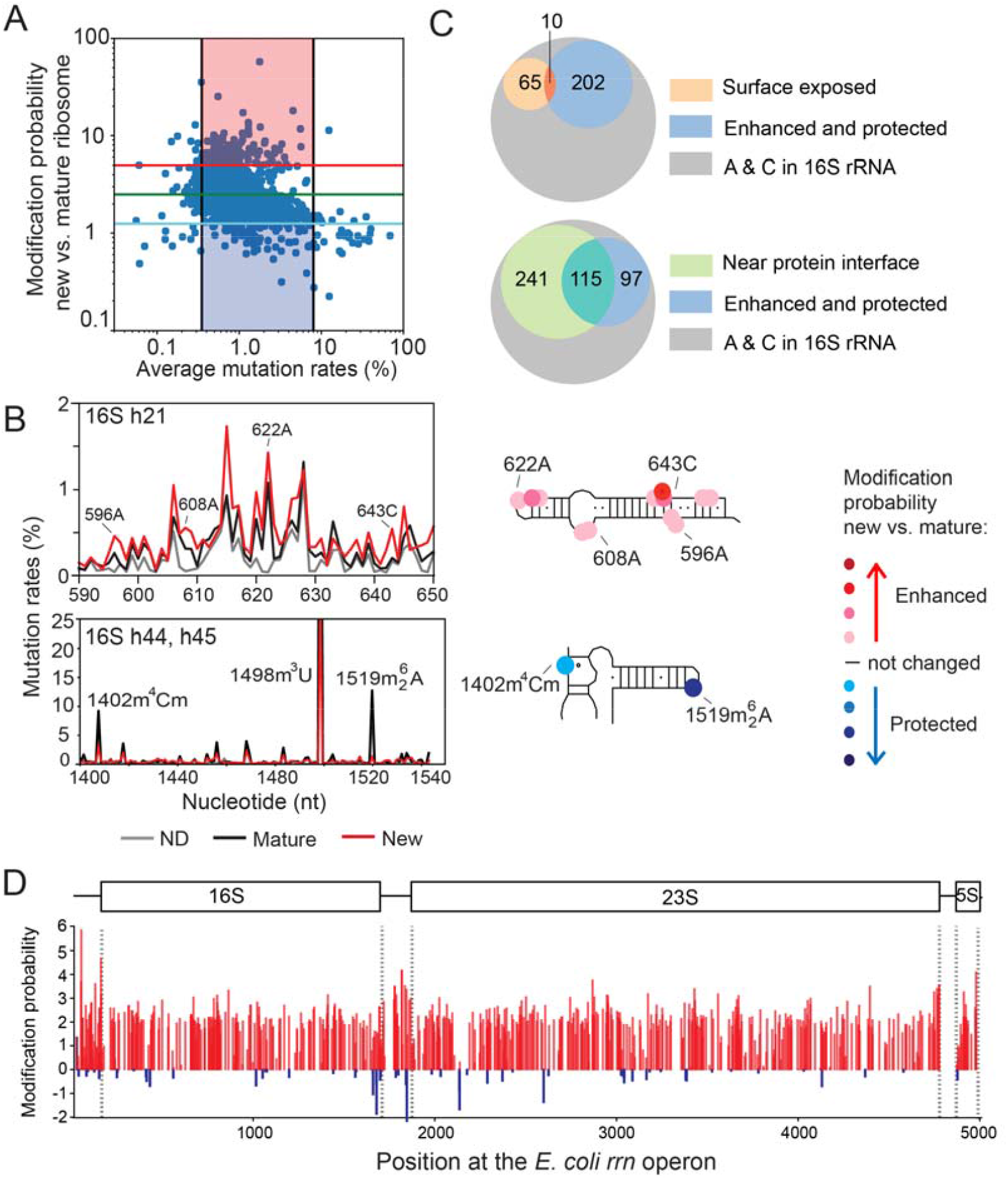
Altered folding of bases throughout the newly transcribed rRNA. A. Sample line plots of mutation rates in 16S h21, 16S h44-45, and 23S H43. Natural modifications in h44 and h45 are labeled. Red, new rRNA 0.5 min after rifampicin; blue, new rRNA after uridine chase; black, mature rRNA; gray, no DMS treatment (ND). B. The relative mutations ratios in new and mature rRNA in 16S h21, 16S h44-45, and 23S H43. The RMR values are calculated from the ratios of mutation rates in (A) as described in Methods. Red and blue lines indicate the high and low cutoffs for enhanced and protected bases, respectively. C. Schematics show RMR for each region of the 16S rRNA, from (B). See Fig. S4 and S5 for further data. 16S U1498 and A1519 appear protected (blue) owing to natural base modification of the mature rRNA. KsgA modifies both A1518 and A1519 (40), but DMS-MaPseq only detected methylation of the downstream A1519 owing to a known difficulty of detecting consecutive A’s by reverse transcription (82,83).

### Comparison of new and mature ribosomal complexes

To map the structural differences between newly made ribosomes and mature ribosomes, we calculated the relative mutation ratios (RMR) for each adenine and cytosine, which is defined as the relative DMS reactivity of that residue in new ^4S^U-labeled rRNA versus the unlabeled mature reference rRNA. A histogram of RMR values showed that many nucleobases were more exposed to DMS in new ribosomes than in mature ribosomes (high RMR), whereas a few bases were protected (low RMR) (Fig. 2, Fig. S2B,C). The same residues were perturbed when ^4S^U incorporation was chased with uridine (Fig. 2B), indicating that these perturbations are not an artifact of rifampicin treatment. A small group of nucleotides with high mutation rates in the untreated ND control corresponded to natural rRNA base modifications that cause misincorporation by TGIRT. Some of these modifications, such as dimethylation of 16S A1518 and A1519 by KsgA (40), occur at a late stage of subunit biogenesis (41,42). A1518 and A1519 were not modified in ^4S^U-labeled rRNA (Fig. 2, Fig. S3B), confirming that our strategy captures immature ribosomal subunits.

RNA from rifampicin-treated cells was systematically two to three-fold more sensitive to DMS modification than the untreated reference (Fig. S2A), which was an effect of the drug and not ^4S^U-labeling (Fig. S2C). This sensitivity to DMS was not specific to rRNA, because the mutation rates of tmRNA were similarly elevated in rifampicin-treated samples. To account for this effect of rifampicin, we benchmarked the DMS probing results against the median value of RMR at 1 min, when the pre-ribosome content was lowest (green line; Fig. 2B). Nucleotides with RMR more than twice or less than half the median were designated ‘exposed’ or ‘protected’ in new ribosomes, respectively (red and blue lines; Fig. 2A). Although pre-rRNA synthesis can initiate at different times during the 1 min ^4S^U-labeling period, potentially blurring structural changes in the rRNA, many bases in new ribosomes showed distinct differences in their susceptibility to DMS modification that were considerably greater than the ND background (Fig. 2B), allowing the broad principles of rRNA folding to be determined.

### Widespread perturbations in new rRNA

To visualize the structural differences between newly made and pre-existing ribosomes, bases that were exposed or protected from DMS modification in new rRNA were mapped onto the mature rRNA secondary and tertiary structures (Fig. 2 and Fig S4-S5). Many A’s and C’s that participate in Watson-Crick base pairs were equally protected from DMS in new and old rRNA within 30 s after rifampicin addition, indicating that much of the rRNA was base paired as expected. Nevertheless, 253 A’s and C’s were more exposed to DMS in new rRNA compared with mature rRNA, indicating that many expected rRNA interactions are not yet in place within this interval.

These conformational differences were distributed across the entire 16S and 23S and 5S rRNAs (Fig. S4B). The absence of any 5′ to 3′ bias up to 2 min after rifampicin addition suggested that stable assembly lags transcription by at least 1 min, which is the time needed to transcribe *E. coli rrn* operons. This apparent lag was initially surprising because stable domains of rRNA tertiary structure can fold within a few seconds (43,44). Moreover, the estimated 2 min total time for subunit biogenesis in *E. coli* had previously suggested that assembly keeps pace with rRNA elongation (2,36). Few of the perturbed residues were on the surface of the 70S ribosomes or the inter-subunit bridges, eliminating subunit joining as an explanation for the differences in DMS modification (Fig. S3). Some of the unfolded bases mapped to regions of the 30S and 50S subunits that are known to assemble slowly, such as the 30S head (16S 3′ domain; Fig. S4) (45,46) or the 50S peripheral stalk base, central protuberance and L1 stalk (Fig. S5) (17). Nevertheless, we also observed structural differences in the 5’ regions of the 16S and 23S rRNAs, which are transcribed first and the most stable *in vitro*. These results suggested that many segments of the rRNA initially form structures that are unstable or heterogeneous.

### Time-dependent folding of rRNA during assembly

To better understand why some interactions in the new rRNA remain partially unfolded, we next asked how the structures of the immature ribosomal complexes change over time. To estimate this, we compared the relative DMS modification (RMR) in samples modified at various intervals after labeling with ^4S^U (Fig. S4, Fig. S5). Those A and C residues that were enhanced or protected compared to the mature rRNA (RMR ≥ 4.6) at more than 1 time-stamped sample were classified according to the pattern of change in RMR over the first 2.5 min after rifampicin treatment (Fig. 3 and Methods). Most nucleotides became more protected by 1 or 1.5 min, consistent with continued assembly and maturation of ^4S^U-labeled complexes (green and orange, Fig. 3). A few residues remained exposed (light red) or became progressively more exposed (red) or more protected (blue) over the entire 2.5 min. These patterns can be explained by interactions that form very late in assembly, or by a stall in the assembly of a subset of complexes in the cell. Finally, certain residues became exposed to DMS at 1.5 min and protected again at 2.5 min (magenta), suggesting opening and closing of the rRNA structure at ∼2 min. As discussed below, nucleotides located near each other in the mature ribosome were protected from DMS with similar kinetics, indicating that the modification patterns reflect coordinated folding events.

**Fig 3.**
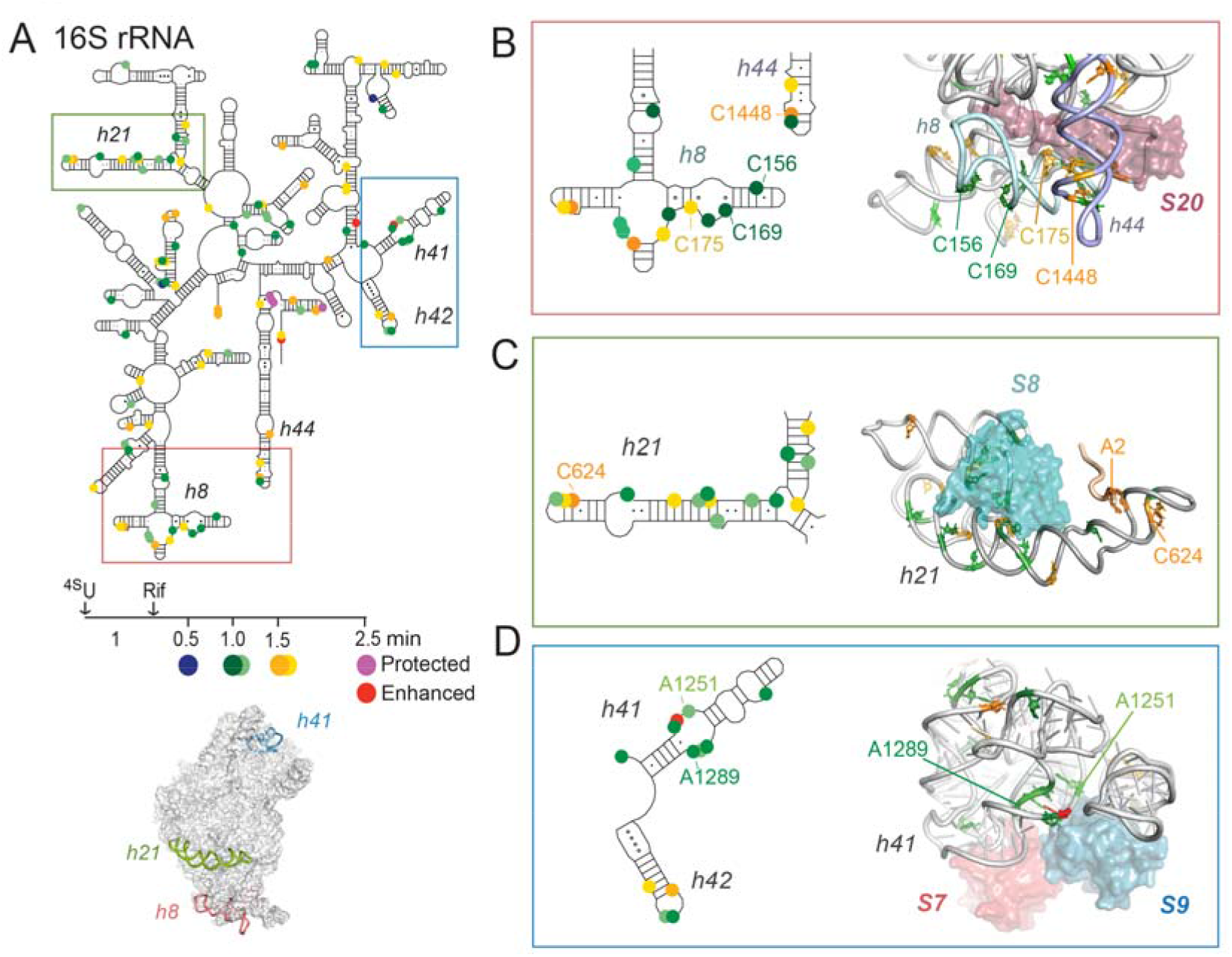
Folding of the rRNA in *E. coli* over time after pulse labeling. Enhanced and protected nucleotides in new ribosomes were clustered based on the direction of change in relative mutation ratio (RMR) in the first 2.5 min after rifampicin addition. See Methods and Fig. S4-S5 for further information. Changes in RMR correlate with increased protection of pre-ribosomes during the initial folding wave (0-1.5 min) and slower conformational changes (≥ 2.5 min). A few bases are increasingly exposed to DMS (red) or temporarily exposed and subsequently reprotected (magenta). Inset plots: Grey lines represent single bases, colored lines indicate the average for each cluster and correspond to the colored circles in the schematics. Open symbols represent bases that were exposed or protected relative to mature ribosomes over the whole 2.5 min. (**A**) 16S rRNA; (**B**) 23S and 5S rRNAs.

### Inter-domain interactions during 30S assembly

Assembly of the 30S subunit has been studied extensively, revealing how r-protein binding to specific helix junctions induces short and long-range re-packing of rRNA helices (11,22). First, 16S helix junctions that are known to switch conformation during 30S assembly *in vitro* exhibited matching time-dependent changes in DMS modification in *E. coli* (Fig. 4A). For example, a 5-helix junction (5WJ) that interacts with proteins uS4 and uS12 (47-49) folded over 1 min (green, Fig. S6A). However, residues that contact uS12 or a flexible region of uS4 were protected more slowly (orange, Fig. S6A).

**Fig 4.**
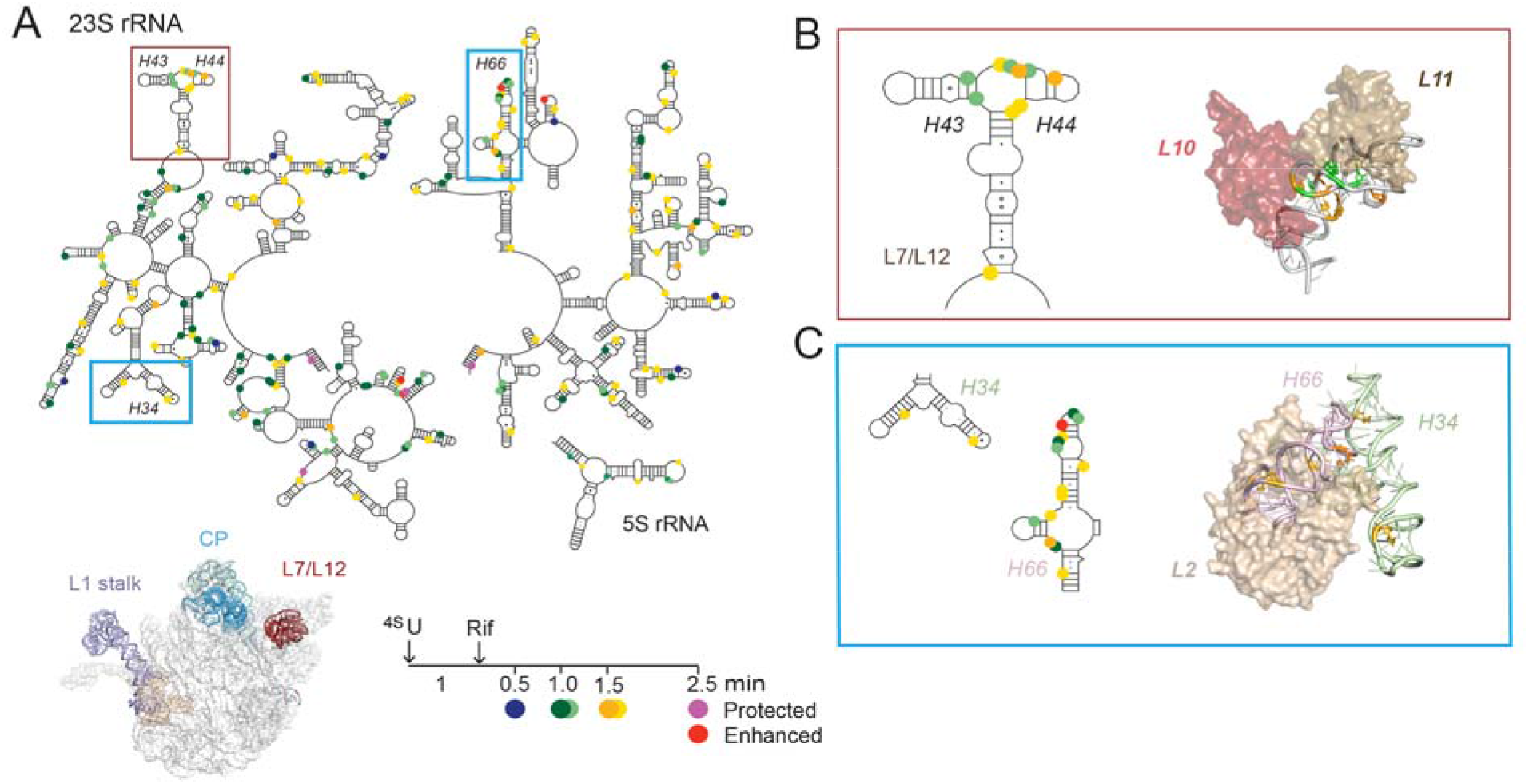
Slow assembly of helix junctions and long-range contacts. Bases that are protected or enhanced over time are colored according to the clusters in Fig. 3. 30S regions of interest are highlighted as colored ribbons in the 16S rRNA and 30S structure (A) and detailed in B-C. 3D ribbons from pdb 4ybb (35). B. 5′ domain: 16S h8 (C193) forms long-range interactions with 16S h44 (C1443, C1448) over 1.5 min. C. 3′ major domain: The internal loop in 16S h41 forms a sharp bend (1 min) at the interface between rRNA subdomains recognized by uS7 and bS9, respectively. The 50S regions of interest (D) that are detailed in E-F. E. A long-range interaction on the subunit interface stabilized by protein uL2. F. Cross-domain helix packing between the 23S H8-H10 region in domain I and H51-H54 in domain III. Residues within H8-H10 fold in 1 min and are stabilized by uL34 (violet); residues that make long-range contacts with H51-54 are protected more slowly. See Fig. S6 for additional details.

Second, we observed slower folding of inter-domain interactions than tertiary interactions within a helix junction. For example, bases that form tertiary interactions around the 16S h8-h10 junction were protected faster than bases that contact the tip of 16S h44 (C193, C1443, C1448) (Fig. 4B), consistent with previous footprinting results (50-52). Similarly, 16S h21 that links protein uS8 in the central domain with bS16 in the 5’ domain (39,53) folded over 1–1.5 min in *E. coli* (Fig. S6B), consistent with time-resolved footprinting of 30S assembly *in vitro* (49,52).

Lastly, we observed the delayed formation of long-range interactions within the 16S 3′ domain (30S ‘head’) that differed from the order in which these helices fold *in vitro. In vitro*, the 30S ‘beak’ (h32-33) and h43 in the uS7 binding site are protected more slowly than other regions of the 16S rRNA (19,54), delaying the recruitment of proteins uS10 and uS14 that are needed to complete the structure of the 30S head (55). This is likely due to misfolding of the nearby helix junctions, which has also been observed during *in vitro* transcription of the 16S 3′ domain (4). By contrast, only a few bases in h32-33 were perturbed in *E. coli* (Fig. 3), suggesting less misfolding in the cell. All the residues of an internal loop in h41 that forms part of the interface between the uS7 and uS9 binding sites was also perturbed in *E. coli* (Fig. 4C), suggesting that h41 and h42 interact after h43 folds. The assembly factor RimM, which acts on 3′ domain h30, h33, and h43 (56,57), may smooth folding of h43 and h33, while delaying tertiary interactions between h41 and h42 (8).

### Slow ordering of peripheral 50S structures

Folding of the 23S rRNA during 50S assembly has been difficult to study because the helical domains are interconnected through many long-range RNA and protein interactions (22), and *in vitro* reconstitution is inefficient (58). However, cryo-EM structures of *in vitro* (17,59) and in-cell assembly intermediates (13,60,61) revealed early organization of a 50S core comprising much of the 23S rRNA, followed by ordering of the central protuberance (CP), L7/L11 stalk base, L1 stalk, and the peptidyl transferase center (PTC) (13,60,61). The importance of core RNA tertiary interactions for 50S assembly was also revealed by pseudouridine interference mapping (62).

In agreement with cryo-EM studies, our in-cell footprinting results showed that the peripheral stalk base, central protuberance, and the L1 stalk remained partially unfolded until 1 to 1.5 min post-rifampicin (Fig. S5). For example, many bases in 23S H43 and H44 that make up the stalk base with proteins uL10 and uL11 and the L1 stalk (Fig. S6C,D) were protected from DMS modification within this time frame. 5S nucleotides that interact with protein uL5 were protected after 1.5 min (Fig. S6E), consistent with reorientation of the CP during 50S and 60S assembly (13,17,63,64).

### Delayed long-range interactions during 50S assembly

In addition to unfolded peripheral structures, we observed slow protection of nucleotides in the core of the mature 50S ribosome that is well-structured in most cryo-EM maps, including 23S domain I which is transcribed first. The distribution of exposed nucleobases throughout the 23S and 5S rRNA suggested that rRNA helices in immature complexes are prone to mis-assembly or partly open to solvent, owing to fluctuations or heterogeneity among the immature complexes.

Some of the slow-folding nucleotides participate in long-range interactions between 23S domains, which are expected to form more slowly than interactions within domains. In the most extreme example, the 5’ and 3’ ends of the 23S rRNA were among the slowest bases to be protected from DMS (magenta, Fig. 3B). These RNA segments are separated by almost 2,900 nucleotides yet need to base pair with each other for pre-23S processing by RNase III. Thus, either this helix has not formed yet in a subset of unprocessed transcripts, or it opens and closes during assembly.

We also observed slow protection of core residues that form long-distance contacts mediated by specific r-proteins. For example, protein uL2 binds H33-35 in 23S domain II while wrapping around 23S H66 in domain IV; these interactions with uL2 form over 1.5 min (orange, Fig. 4E). The helices recognized by uL2 pack against two other helix junctions that form another long-range contact: bL34 and H8-10 in 23S domain I (green, 1 min folding time) and H51-54 in 23S domain III (orange, 1.5 min folding time; Fig. 4F). Although each helix junction folds within the same time window as other tertiary interactions in the 50S ribosome (green and orange, 1-1.5 min), bases on the interface between H8-10 and H65-66 them take ≥ 2.5 min to become fully protected (red, magenta; Fig. 4E-F), suggesting that this region remains flexible until the end of 50S assembly. Interestingly, this set of long-range interactions connecting 23S domains I, II, III and IV sandwiches the base of the L1 stalk, and its integrity is likely essential for 50S function. The complexity of these long-range interactions suggests that they could form in more than one temporal order, and indeed, uL2 was proposed to join the complex at different stages depending on the path of 50S assembly (13).

### Active site rearrangements occur late in assembly

In the cell, assembly factors assist folding of the active sites, while acting as quality control checkpoints to ensure that the new ribosomes are competent for translation (22,23). During *in vitro* reconstitution or in cells deficient for assembly factors or r-proteins, the active sites for peptide synthesis and decoding are the last regions to fold (8,17,46,52,65). In our results, however, the 23S PTC, 16S decoding site (h44) and central pseudoknot (h2) were more protected from DMS in new ribosomes than in mature ribosomes (neutral or blue symbols, Fig. S4 and S5). This strong protection suggested that the active sites in immature complexes are continually shielded by assembly factors in *E. coli*. It could also arise from misfolding of the active sites, although the protection is more extensive than would be expected for local mispairing.

Although most active site nucleobases were protected from DMS, we observed distinct signatures of conformational change in the 16S and 23S rRNA around the tRNA P-site and mRNA binding platform that occur after other interactions in the 30S and 50S subunits have been established. These changes are consistent with remodeling of the active sites during the binding and release of assembly factors. In the 16S rRNA, residues in h44 and h45 were exposed from 0.5 – 1.5 min and then protected from 1.5 – 2.5 min, consistent with refolding of this region at a late stage of 30S assembly (magenta, Fig. 5A). In the mature 30S subunit, these nucleotides form a tertiary contact between h44 and the loop of h45. This contact is dislocated by binding of the assembly factor RsgA to immature subunits and then repositioned after release of RsgA and binding of initiation factor IF3 and initiator tRNA (9,66-70). We also observed that the loop of h45 was not yet methylated by KsgA (Fig. 2C), which is thought to be one of the latest steps in 30S maturation (41,71). By contrast, 16S U1498 was fully methylated by RsmE (72), and 16S C1402 was partially methylated by RsmH and RsmI (73) in new ribosomes (Fig. 2C), consistent with the idea that these enzymes act earlier in 30S biogenesis.

**Fig 5.**
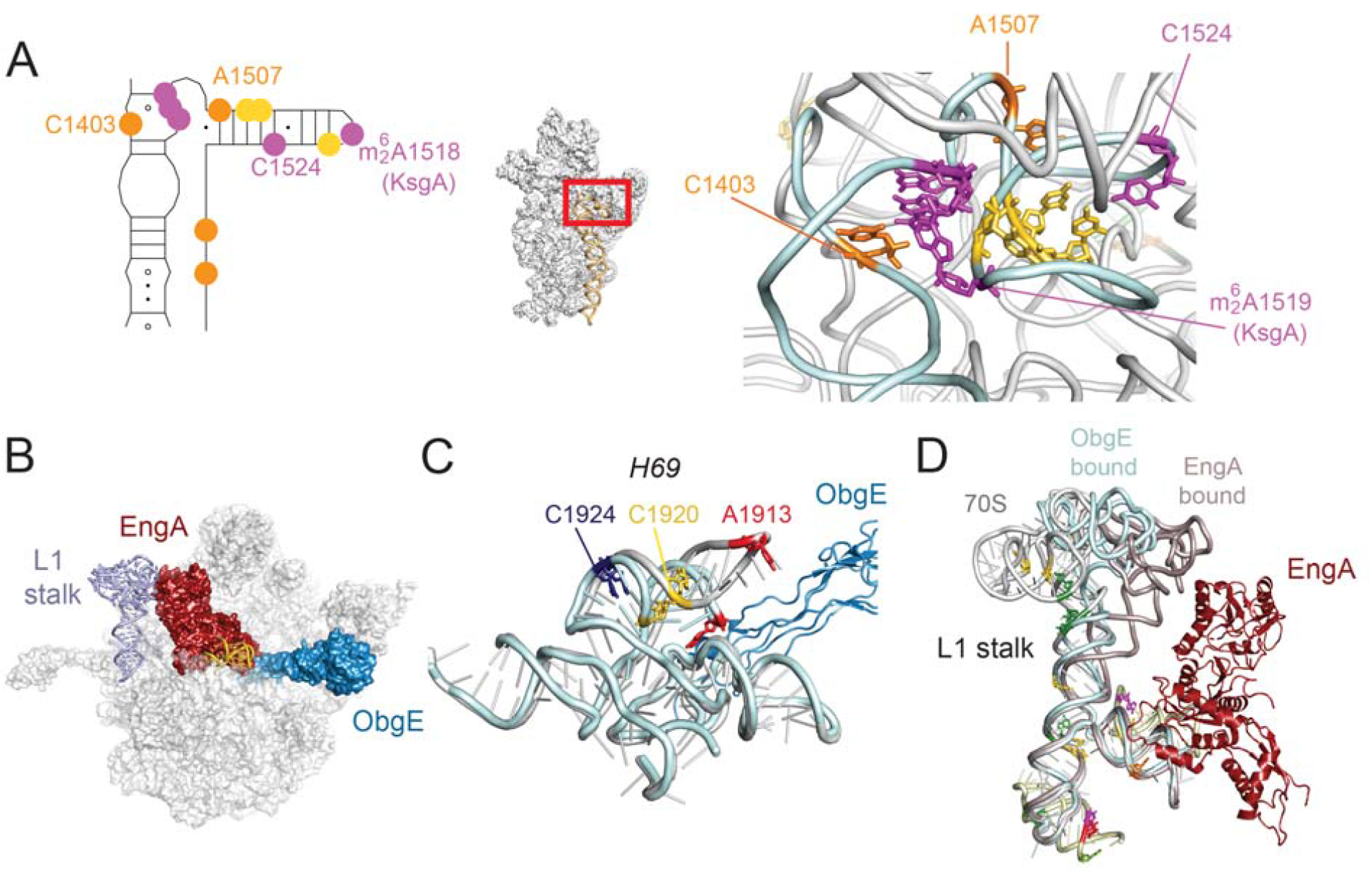
Late-stage remodeling of tRNA binding sites in 30S and 50S subunits. A. 30S mRNA binding platform and decoding site formed by h44 and h45 (wheat and cyan), with enhanced and protected residues colored by the folding kinetics as in Fig. 3. Nucleotides in magenta are exposed to DMS at 1-1.5 min and reburied by 2.5 min, indicating a conformational change. Natural rRNA modification sites are also shown. B. Superposition of 50S subunits in complex with assembly factors, showing how ObgE (blue; pdb 4csu (76)) and EngA (maroon; pdb 3j8g (75)) interact with the L1 stalk (light blue) and H69 (gold). C. 23S nucleotides that undergo late-stage structural rearrangements (magenta) include residues below the L1 stalk. Superposition of the L1 stalk in 70S (grey), ObgE bound-50S (pale cyan), and EngA-bound 50S ribosomes (aluminum). D. Enhanced modification of A1913 in H69 correlates with a transition from a pre-50S conformation bound by ObgE (pale cyan) to a mature 50S structure (grey; pdb 4ybb).

In the 23S rRNA, we observed a similar pattern of active site protection (Fig. S5B) and late remodeling of H69 and the L1 stalk that is consistent with the release of 50S assembly factors. Although the L1 stalk moves with the E-site tRNA during translation (74), it adopts even more extreme orientations when in complex with assembly factors ObgE and EngA (75,76) (Fig. 5B,C) or YjgA (14). Consistent with the idea that the L1 stalk must also flex during 50S maturation, we observed that residues surrounding the base of the L1 stalk were exposed to DMS and refolded around 2.5 min (magenta; Fig. 5C). 23S H69 extends into the decoding site of the 70S ribosome, forming the intersubunit bridge B2a. A1913 at the tip of H69 was protected at 0.5 min post-rifampicin but became more exposed over our 2.5 min time course (red; Fig. 5D). In free 50S subunits, H69 projects into the solvent (77), explaining why A1913 becomes more exposed to DMS as 50S subunits mature. By contrast, a cryo-EM structure of the 50S ribosome bound to the assembly factor ObgE shows H69 rotated inward (Fig. 5D), with the Watson-Crick edge of A1913 buried in a kink in the adjacent helix (76). In a more recent cryo-EM structure of a ObgE-bound pre-50S complex, the pseudouridine synthase RluD covers the tip of H69, which may also explain why A1913 is protected in the immature complexes (14). Thus, for both 30S and 50S ribosomes, specific conformational changes near the assembly factor binding sites occurred later than other steps of assembly.

## DISCUSSION

### Time-resolved DMS footprinting of ribosome assembly in *E. coli*

The biosynthesis and assembly of ribosomes requires only a few minutes in rapidly dividing bacteria, making it challenging to follow the assembly process in real time. In this study, we show that metabolic pulse-labeling of pre-rRNA transcripts with 4-thiouridine and rifampicin, combined with high-throughput DMS footprinting (4U-DMS-MaP), captures structural changes in newly made rRNA in *E. coli* cells. The advantage of this approach is that the rRNA structure is probed *in situ* during normal growth, without a need for mutations that stall or divert assembly. Although the 1 min labeling time means that the transcripts pulled down by ^4S^U are present in a mixture of complexes, this approach still captures the major stages of rRNA folding during assembly. We cannot exclude the possibility that incorporation of ^4S^U slows maturation of the pre-30S and pre-50S particles. However, labeled and unlabeled mature complexes have the same DMS modification profile, indicating that any perturbation is moderate under our conditions.

### Slow consolidation of tertiary interactions after transcription

Our DMS footprinting results showed that the initial ribosomal complexes are more dynamic or heterogeneously folded than previously supposed (Fig. 6). Although the rRNA secondary structure is expected to base pair within the 1 min time scale of the in-cell footprinting experiment, a surprising number of bases remain exposed to DMS modification long after transcription is done. Many of these bases participate in rRNA tertiary interactions or protein interactions in the mature ribosome. These perturbations were spread across the operon, including the 16S 5’ domain and 23S domain I, which are the first to be transcribed and typically ordered in single particles (17). This evidence for wide-spread perturbations to the rRNA interactions suggests that helix junctions may fold locally soon after the rRNA is transcribed, but the overall structure of the particle is not finalized until larger domains can be consolidated (Fig. 6). This result agrees with recent findings that individual proteins are not stably recruited to the pre-rRNA until multiple r-proteins bind the transcript (3,4).

**Fig. 6.**
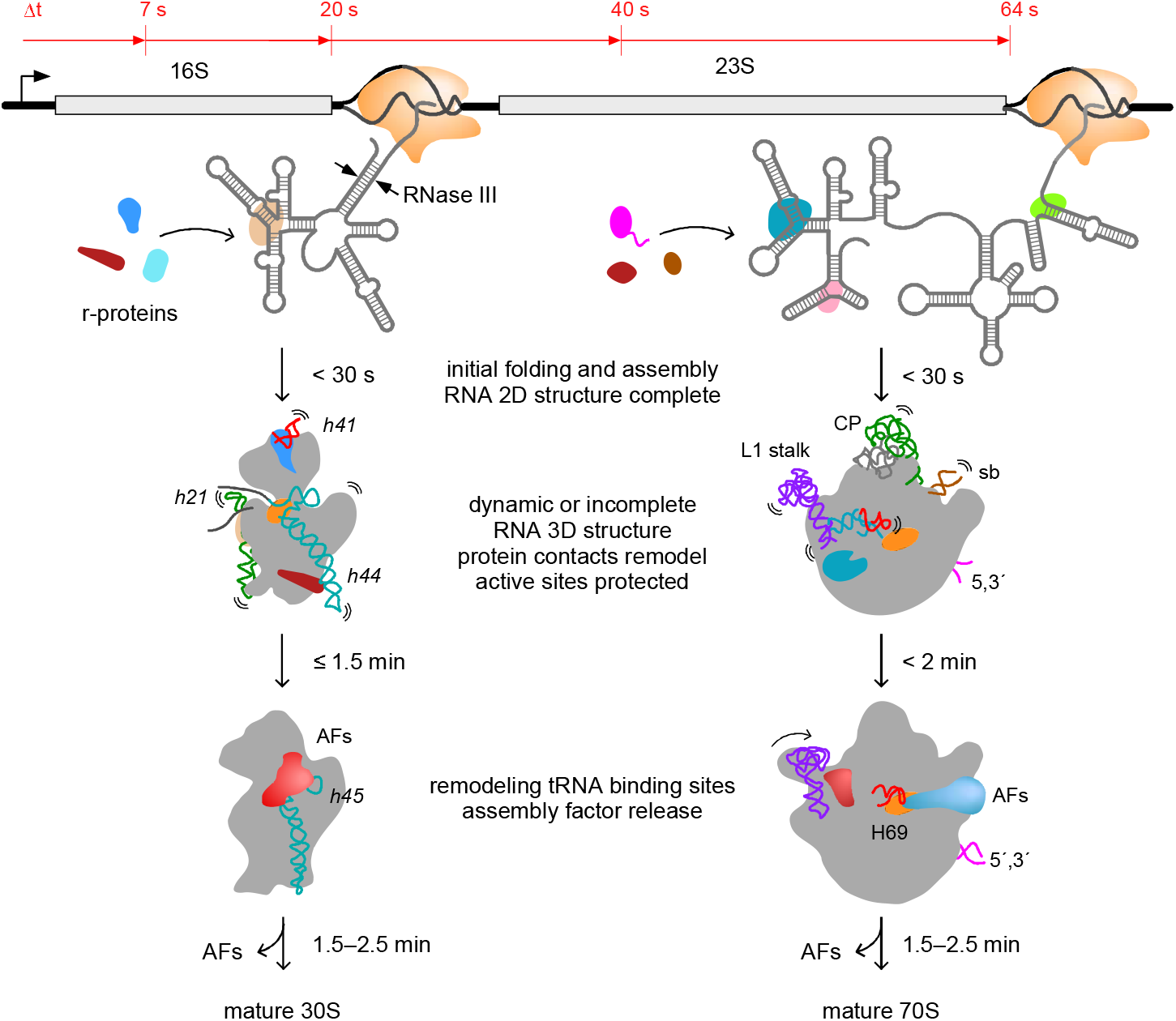
Models of rRNA folding during ribosome assembly in *E. coli*. 4U-DMS-MaP reveals details of rRNA conformation and assembly upon pre-rRNA synthesis in growing *E. coli* cells. Top, average time line for rRNA transcription, from (84). Bottom, key stages of ribosome assembly inferred from these results and cryo-EM studies (see text). Although rRNA bases are expected to pair soon after transcription, rRNA and r-protein interactions evolve over 1–2 minutes in an ensemble of pre-ribosomal complexes that remain open to DMS modification. Assembly is followed by structural rearrangements in 16S h45, 23S L1 stalk and 23S H69 that correlate with assembly factor (AF) binding and release. The 16S central pseudoknot and A-site and the 50S peptidyl transferase center remain protected during assembly, presumably by additional AFs (orange). Grey outlines cartooned from pdb 2ykr (69), 6gc4, 6gc6 (17), 4csu (76), 3j8g (75), YjeQ bound cryoEM structures and Era depleted cryo-EM structures (5,65).

We frequently observe that long-range tertiary interactions fold more slowly than short-range tertiary interactions, as previously seen in time-resolved footprinting of 30S assembly *in vitro* (52). Our in-cell footprinting shows that this delay is even more common in the 50S subunit, which contains many interlocking structural modules that are far apart in sequence. Slow formation of long-range tertiary interactions is expected during transcription, owing to the time needed to synthesize downstream interacting partners. In agreement with the idea that transcription contributes to such delays, base pairing between the 5’ and 3’ ends of the 23S rRNA, which takes ∼30 s to transcribe, was one of the slowest folding events that we detected. Importantly, our results suggest that the delay in long-range interactions also renders local structures unstable or partly unstable.

### Reorganization of primary protein binding sites

*In vitro* experiments have shown that the rRNA and r-proteins change structure after they come together (39,47,52), which is important for the cooperativity of assembly (78-80). Counterintuitively, time-dependent changes in DMS reactivity were more prevalent in the binding sites of primary 30S assembly proteins than of late assembly proteins in the Nomura map (Fig. S7). One possibility is that the binding sites of primary assembly proteins need longer times to fold correctly, limiting the overall speed of assembly. Alternatively, these primary rRNA-protein interactions may remodel as assembly progresses, whereas late assembly proteins may have little further influence on the structure of the rRNA (39).

### Shielded active sites during late biogenesis

Late ribosome assembly factors in bacteria and yeast occupy active sites, presumably protecting these vulnerable regions of the rRNA from nucleases, while preventing immature large and small subunits from engaging in translation (1). Remarkably, we find that the PTC and decoding center are more protected from DMS on average than the reference mature ribosomes throughout the time course of our experiment, even when peripheral regions of each subunit are partially unfolded. By contrast, earlier footprinting experiments showed that nucleotides surrounding the active sites were highly exposed to solvent in *E. coli* lacking a specific assembly factor (8,65). Altogether, these observations suggest that the active site regions do not self-assemble efficiently in the cell and must be continually shielded by assembly factors that may also assist their folding.

Although most active site residues are protected from DMS, a few nucleotides change conformation after the tertiary structure and rRNA-protein interactions have formed (Fig. 6). The change in DMS accessibility correlates well with perturbations from the binding and release of assembly factors revealed by cryo-EM structures of ribosomal complexes bound to assembly factors. Candidates include RbfA and RsgA (YjeQ) that interact with 16S h44 and h45, EngA (Der) that interacts with the 50S L1 stalk, and ObgE and the RluD synthase that alters the conformation of 23S H69. These refolding events occur after the other regions of the rRNA have assembled and are clustered around the P-site and mRNA binding site, consistent with their proposed role in the quality control of ribosome assembly.

### Future applications of 4U-DMS-MaP

DMS modification is particularly useful for probing conformational changes in regions of the rRNA that are too flexible or too fragile to be visible by cryo-EM. In a recent study of Era GTPase, for example, helices that were missing from cryo-EM structures of the 30S complexes were shown by DMS to be mainly base paired, suggesting some limitations of cryo-EM in detecting dynamic RNA structures (65). Since 4U-DMS-MaP probes the rRNA in its natural environment and under biological time scales, our results provide new insight into the process of ribosome assembly as it occurs in actively growing cells.

Since ^4S^U incorporates into all new transcripts during the labeling time window, the method can be used to study other structured RNAs and RNA-protein complexes in *E. coli* cells. Because metabolic labeling with ^4S^U and other base analogs is widely used to study transcriptomes in yeast and mammalian cells (30,32), our 4U-DMS-MaP method can be adapted to study RNA structures in other organisms, as long as the transcripts are sufficiently numerous to detect after a brief labeling period.

One challenge of in-cell RNA footprinting is the conformational heterogeneity of newly synthesized transcripts. Although the DMS modification pattern represents the population average, it is notable that we still observe distinct patterns reflecting the slowest folding events in the cell. The ability to resolve different assembling complexes depends on the time resolution of pulse labeling, which is currently limited by ^4S^U permeability and incorporation efficiency. The method can be improved in the future by achieving shorter pulse labeling times or through fractionation of transcription complexes with affinity tags (81).

## Acknowledgements

The authors thank the Johns Hopkins Genetic Resources Core Facility high-throughput sequencing core for technical support. The authors thank Dr. Yiliang Ding, Dr. Sander Granneman and all members of the Woodson lab for helpful discussions.

## RESOURCE AVAILABILITY

### Materials & Correspondence

Further information and requests for reagents may be directed to Sarah Woodson.

### Data and Code Availability

All software and algorithms are listed in Table S1. The accession number of 4U-DMS-MaP sequencing datasets is GEO: GSE157014. The analysis pipeline can be found on Github: https://github.com/woodsonlab/4U-DMS-MaP.

