## Supplemental Information for "Late consolidation of rRNA structure during co-transcriptional assembly in *E. coli* by time-resolved DMS footprinting"

The sample treated with DMS 1 min after rifampicin had the lowest median RMR and represented the state that was the closest to the mature ribosome. The median RMR of this 1 min post-rifampicin sample, $\overline{RMR}_{\text{Rif1}}$, was used to benchmark the perturbations to new ribosomes for the entire time course. Nucleotides with $RMR_i > 2 \times \overline{RMR}_{\text{Rif1}}$ were designated 'enhanced', and those with $RMR_i < 0.5 \times \overline{RMR}_{\text{Rif1}}$ were designated 'protected'. The enhanced and protected nucleotides were mapped on the rRNA secondary structures with xRNA and 3D structures with PyMOL.

**Funding:** This work was supported by grants from the National Institutes of Health (R35GM136351, R01NS099397 to S.A.W.)

**Competing interests:** The authors declare no competing interests.

**RESOURCE AVAILABILITY**

**Data and Code Availability:** All software and algorithms are listed in Table S1. The accession number of 4U-DMS-MaP sequencing datasets is GEO: GSE157014. The analysis pipeline can be found on Github: https://github.com/woodsonlab/4U-DMS-MaP.

**FIGURES AND FIGURE LEGENDS**

Fig. 1

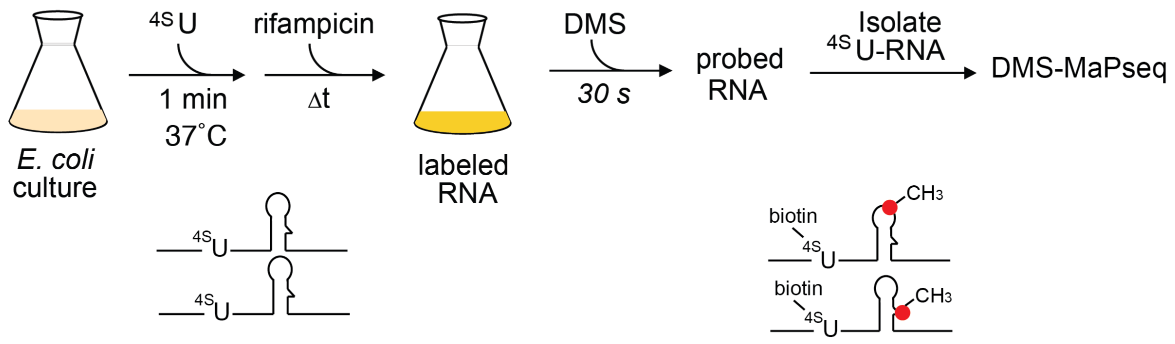


**Fig 1. Visualizing the structures of newly made ribosomes with 4U-DMS-MaP.**

A. Overview of metabolic labeling of new transcripts and DMS footprinting in E. coli. Labeling with 4-thiouridine was stopped by rifampicin to inhibit transcription initiation or chased with excess uridine; 1 min $^{4S}$U labeling and 30 s of DMS treatment were optimal for high-throughput footprinting of new rRNA (Fig. S1A).

B. Assembly kinetics based on the amount of $^{3}$H-uridine in peak fractions of sucrose gradient polysome profiles (Fig. S1D, H). The RNA was labeled with $^{3}$H-uridine for 1 min at 37 ºC before rifampicin addition (solid lines), or for 2 min at 25 ºC before uridine addition (dashed lines). The $^{3}$H-U content in the 30S and 50S peaks are plotted in red and blue, respectively. The labeled subunits accumulate about twice as slowly at 25 ˚C than at 37 ˚C.

Fig. 2

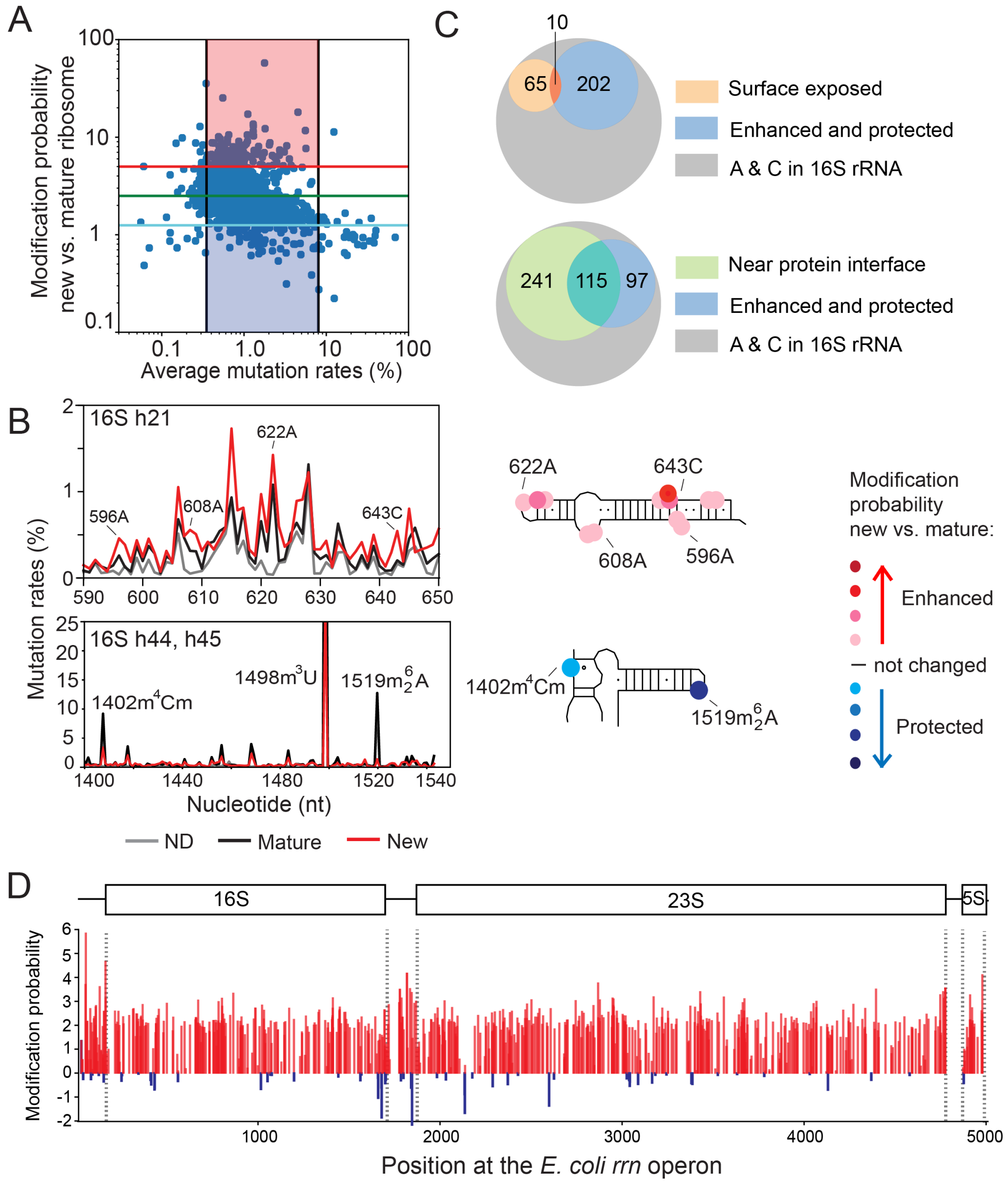


**Fig 2. Altered folding of bases throughout the newly transcribed rRNA.**

A. Sample line plots of mutation rates in 16S h21, 16S h44-45, and 23S H43. Natural modifications in h44 and h45 are labeled. Red, new rRNA 0.5 min after rifampicin; blue, new rRNA after uridine chase; black, mature rRNA; gray, no DMS treatment (ND).

B. The relative mutations ratios in new and mature rRNA in 16S h21, 16S h44-45, and 23S H43. The RMR values are calculated from the ratios of mutation rates in (A) as described in Methods.

Red and blue lines indicate the high and low cutoffs for enhanced and protected bases, respectively. C. Schematics show RMR for each region of the 16S rRNA, from (B). See Fig. S4 and S5 for further data. 16S U1498 and A1519 appear protected (blue) owing to natural base modification of the mature rRNA. KsgA modifies both A1518 and A1519 (40), but DMS-MaPseq only detected methylation of the downstream A1519 owing to a known difficulty of detecting consecutive A's by reverse transcription (82,83).

Fig. 3

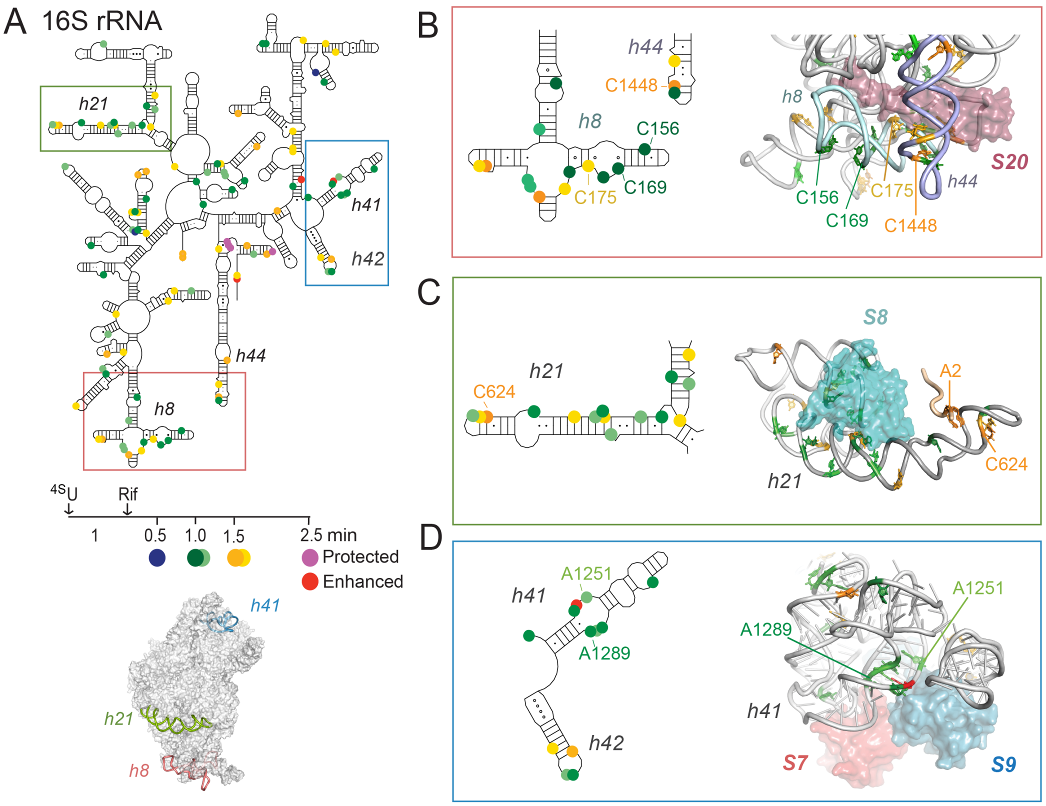


**Fig 3. Folding of the rRNA in *E. coli* over time after pulse labeling.**

Enhanced and protected nucleotides in new ribosomes were clustered based on the direction of change in relative mutation ratio (RMR) in the first 2.5 min after rifampicin addition. See Methods and Fig. S4-S5 for further information. Changes in RMR correlate with increased protection of pre-ribosomes during the initial folding wave (0-1.5 min) and slower conformational changes (≥ 2.5 min). A few bases are increasingly exposed to DMS (red) or temporarily exposed and subsequently reprotected (magenta). Inset plots: Grey lines represent single bases, colored lines indicate the average for each cluster and correspond to the colored circles in the schematics. Open symbols represent bases that were exposed or protected relative to mature ribosomes over the whole 2.5 min. (**A**) 16S rRNA; (**B**) 23S and 5S rRNAs.

Fig. 4

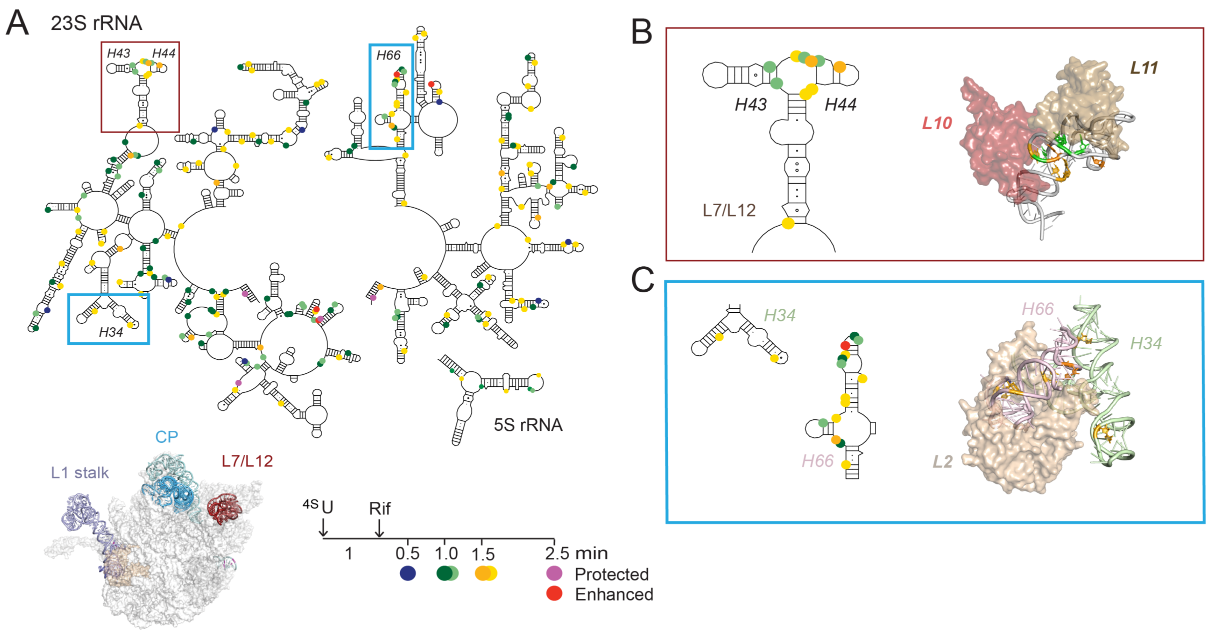


**Fig 4. Slow assembly of helix junctions and long-range contacts.**

Bases that are protected or enhanced over time are colored according to the clusters in Fig. 3. 30S regions of interest are highlighted as colored ribbons in the 16S rRNA and 30S structure (A) and detailed in B-C. 3D ribbons from pdb 4ybb (35). B. 5′ domain: 16S h8 (C193) forms long-range interactions with 16S h44 (C1443, C1448) over 1.5 min. C. 3′ major domain: The internal loop in 16S h41 forms a sharp bend (1 min) at the interface between rRNA subdomains recognized by uS7 and bS9, respectively. The 50S regions of interest (D) that are detailed in E-F. E. A long-range interaction on the subunit interface stabilized by protein uL2. F. Cross-domain helix packing between the 23S H8-H10 region in domain I and H51-H54 in domain III. Residues within H8-H10 fold in 1 min and are stabilized by uL34 (violet); residues that make long-range contacts with H51-54 are protected more slowly. See Fig. S6 for additional details.

Fig. 5

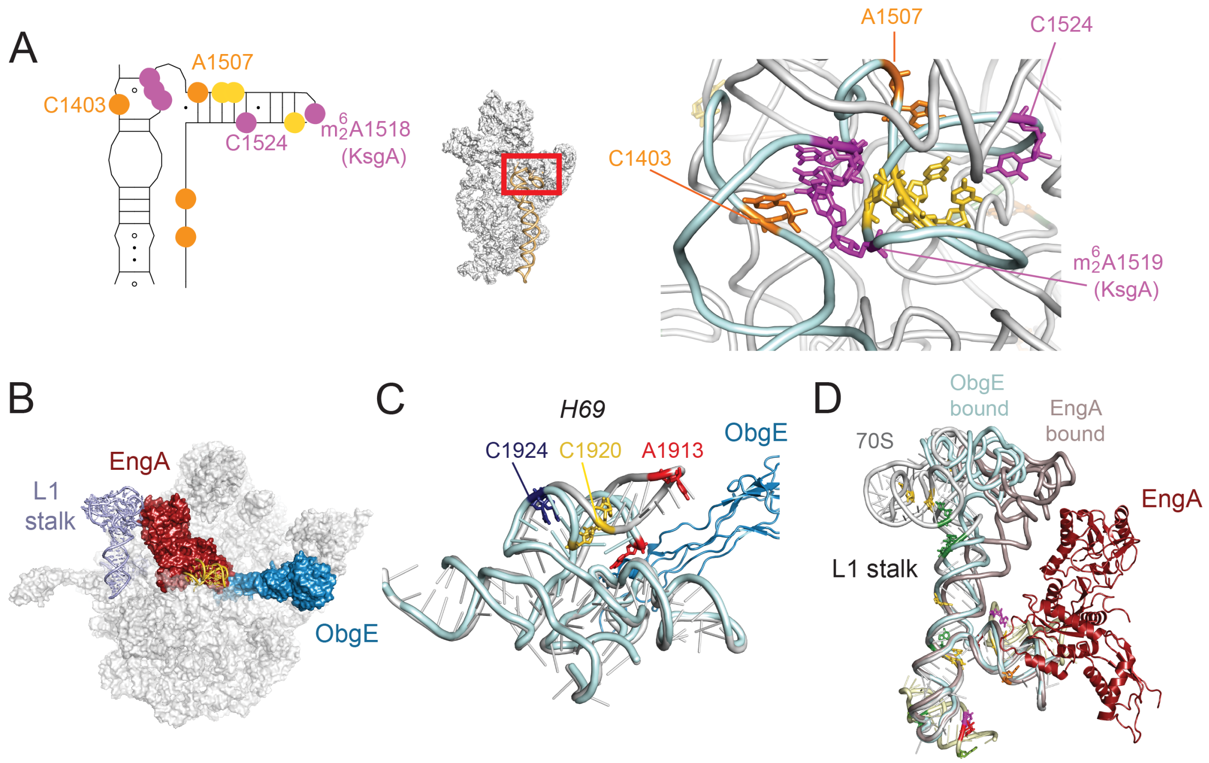


**Fig 5. Late-stage remodeling of tRNA binding sites in 30S and 50S subunits.**

A. 30S mRNA binding platform and decoding site formed by h44 and h45 (wheat and cyan), with enhanced and protected residues colored by the folding kinetics as in Fig. 3. Nucleotides in magenta are exposed to DMS at 1-1.5 min and reburied by 2.5 min, indicating a conformational change. Natural rRNA modification sites are also shown. B. Superposition of 50S subunits in complex with assembly factors, showing how ObgE (blue; pdb 4csu (76)) and EngA (maroon; pdb 3j8g (75)) interact with the L1 stalk (light blue) and H69 (gold). C. 23S nucleotides that undergo late-stage structural rearrangements (magenta) include residues below the L1 stalk. Superposition of the L1 stalk in 70S (grey), ObgE bound-50S (pale cyan), and EngA-bound 50S ribosomes (aluminum). D. Enhanced modification of A1913 in H69 correlates with a transition from a pre-50S conformation bound by ObgE (pale cyan) to a mature 50S structure (grey; pdb 4ybb).

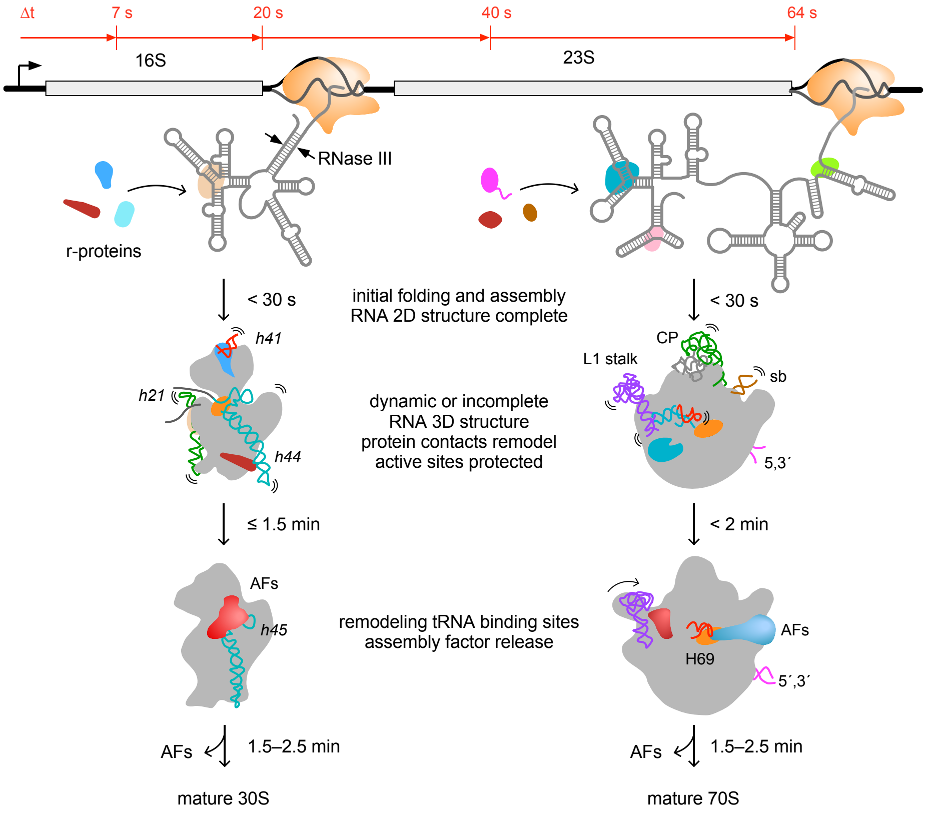


**Fig. 6. Models of rRNA folding during ribosome assembly in *E. coli*.**

4U-DMS-MaP reveals details of rRNA conformation and assembly upon pre-rRNA synthesis in growing *E. coli* cells. Top, average time line for rRNA transcription, from (84). Bottom, key stages of ribosome assembly inferred from these results and cryo-EM studies (see text). Although rRNA bases are expected to pair soon after transcription, rRNA and r-protein interactions evolve over 1–2 minutes in an ensemble of pre-ribosomal complexes that remain open to DMS modification. Assembly is followed by structural rearrangements in 16S h45, 23S L1 stalk and 23S H69 that correlate with assembly factor (AF) binding and release. The 16S central pseudoknot and A-site and the 50S peptidyl transferase center remain protected during assembly, presumably by additional AFs (orange). Grey outlines cartooned from pdb 2ykr (69), 6gc4, 6gc6 (17), 4csu (76), 3j8g (75), YjeQ bound cryoEM structures and Era depleted cryo-EM structures (5,65).
